# Postural and muscular effects of upper-limb movements on voicing

**DOI:** 10.1101/2023.03.08.531710

**Authors:** Wim Pouw, Lara S. Burchardt, Luc Selen

## Abstract

Voice production can be a whole-body affair: Upper limb movements physically impact the voice in steady-state vocalization, speaking, and singing. This is supposedly due to biomechanical impulses on the chest-wall, affecting subglottal pressure. Unveiling such biomechanics is important, as humans gesture with their hands in a synchronized way with speaking. Here we assess biomechanical interactions between arm movements and the voice, by measurement of key (respiratory-related) muscles with electromyography (EMG) during different types of upper limb movement while measuring the bodys center of mass. We show that gesture-related muscle activations scale with positive peaks in the voices amplitude. Some of these muscles also strongly associate with changes in the center mass, confirming that gesture-vocal coupling partly arises due to posture-related muscle activity. If replicated, these results suggest an evolutionary ancient gesture-vocal connection at the level of biomechanics. These preliminary results will support a pre-registration of analyses for a larger-scale confirmatory study.

## INTRODUCTION

In principle, any muscle that attaches to the rib cage can act on it and thus affect the subglottal pressure that supports voicing. Consequently, there are a lot of potential respiratory-vocal muscles. Said muscles would include those around the chest (e.g., pectoralis major), abdomen (rectus abdominus), and back (erector spinae, seratus posterior/anterior). However, in passive breathing and speaking, only a subset of the possible muscles are used. Most notably, the diaphragm and the muscles between the ribs (intercostalis) drive passive speech-supporting respiration ^1,17^. Only on rarer occasions, when coughing, shouting, or breathing deeply, humans recruit other so-called accessory respiratory muscles such as the abs and pectoral muscles ^1,7,17^. From this it is tempting to conclude that there is only a small set of primary respiratory muscles.

Yet, when humans speak or sing it is common to move the upper limbs expressively at the same time called gesture^11,20^. Such upper limb movements will recruit a whole range of upper body muscles, including those involved in maintaining posture^4^. Several of these muscles attach to the rib cage (e.g., ab- and pectoral muscles). These muscles are classically listed as accessory to respiratory functioning^17^. Given that speaking requires subtle modulations of subglottal pressure^16,18,19^, co-gesture speaking must be in some way coordinated with the respiratory-and-thus-vocal muscles that are activated during gesturing (for an overview see^12^).

There is now considerable evidence that gestural arm movements affect the voice directly,as summarized in Pouw Fuchs (2022)^12^: More extreme peaks in the acceleration of movements with higher (vs. lower) mass upper limb segments relate to changes in chest circumference^14^, which associated with more extreme acoustic effects on the intensity of vocal sound (through increasing subglottal pressure). Furthermore, acoustic effects of upper limb movements are more extreme when subjects are in a more unstable standing versus sitting position^13^. These previous studies assessed continuous voicing, mono-syllable utterances, and fluent speech production^15^. This ties in with the idea that a physical impulse (mass x acceleration), impacts posture (especially when standing), recruiting respiratory-related muscles (that change chest circumference), which impacts respiratory-vocal functioning (such that intensity and F0 are affected). Such interactions, between pectoral/upper limb activity and respiratory-vocal states, have been well-studied in non-human animals^2,3,6^.

Participants perform no movements (see here) as well as 4 actions along the sagittal (flexion [see here], and extension, [see here]) and transversal plane (internal [see here], and external rotation [see here]) with the elbow flexed at a 90-degree angle. During these movements, they produce a steady-state schwa-like phonation with one egressive flow. As we know that gesture-speech biomechanics is modulated by posture ^13^ and some of those postural muscles are well-known as expiratory muscles, we measure two postural related muscles: the rectus abdominus, and the erector spinae in the back. Both the abs and erector spinae directly attach to the rib cage. We also assess focal muscles with and without well-known respiratory effects: We measure an internal rotator muscle on the chest (pectoralis major), and an external rotator muscle on the scapula (infraspinatus). The pectoralis major is associated with respiratory interactions as reviewed above and directly attaches to the rib cage. The infraspinatus is not directly attached to the rib-cage, but it does insert into the scapula which is a dynamic stabilizer for the entire should girdle region ^8^. Thus, even indirectly it might have a role to play in gesture-speech biomechanics, but most likely, weakly so. See figure 1 for the muscles measured with sEMG in this study.

**Figure 1:**
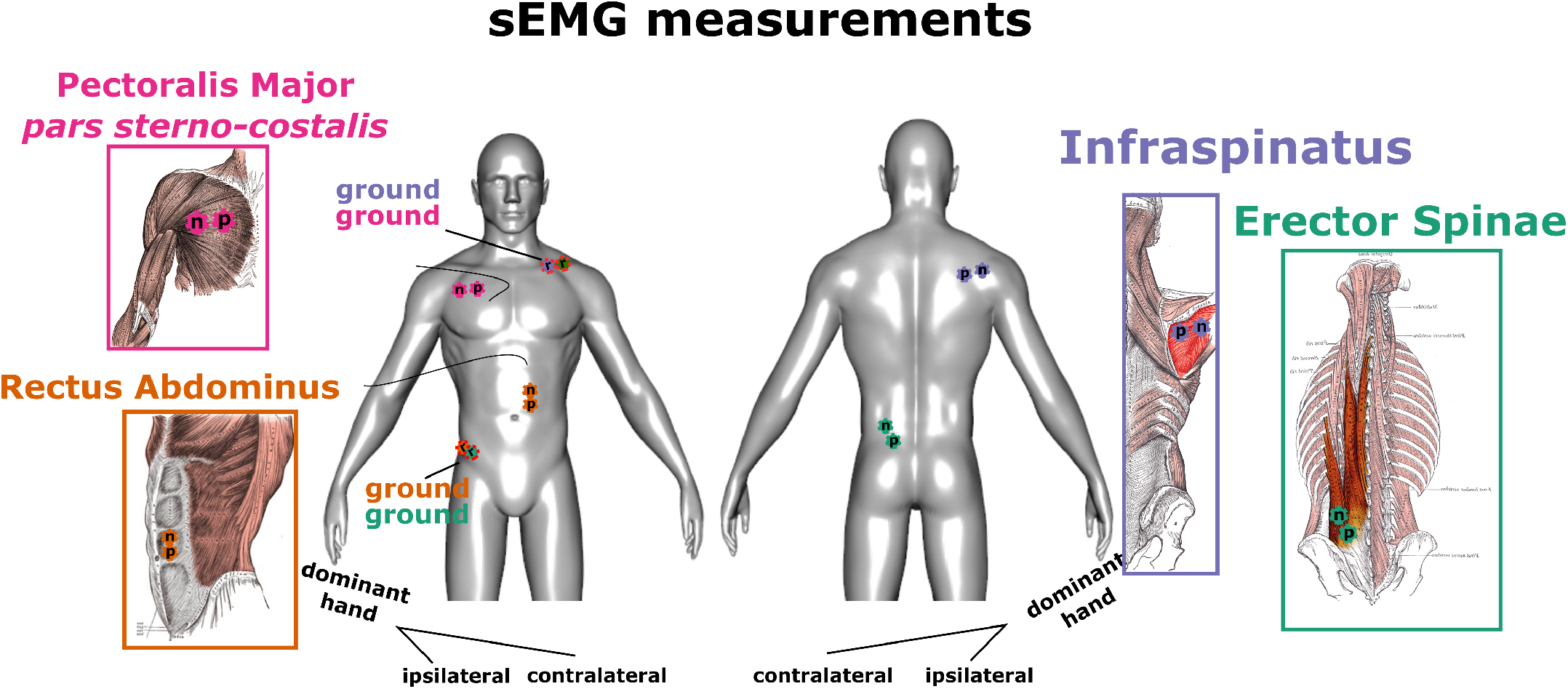
Muscle measurements performed for this study.

## METHODS

The current study has been approved by the Ethics Committee Social Sciences (ECSS) of the Radboud University, reference nr.: 22N.002642. The current study serves as a preliminary study for a larger planned study of over 15 individuals.

### Participants

For the current pilot experiment, the first author (Dutch-speaking; male; right-handed; age 35; 21.7 BMI; 68 kg; length 173cm; upper arm circumference = 31.5cm; triceps skinfold = 16mm) and a volunteer female (Dutch-speaking; right-handed; age 37; 57 kg; 21.5 BMI; length 161cm; upper arm circumference = 26cm; triceps skinfold = 21mm) performed the experiment.

### Study design

The 1-hour study involves a two-level within subject factor wrist-weight manipulation (no weight vs. weight), a two-level within subject vocalization condition (expire vs. vocalize), and a five-level within-subject movement condition (no movement, extension, flexion, external rotation, internal rotation). With 4 trial repetitions over the experiment, we yield 80 trials per participant. Trials were blocked by weight condition and vocalization condition (so that weights and tasks did not switch from trial to trial). Within blocks, all movement conditions were randomized in order of presentation.

### Measurements and equipment

#### Weight wrist

To manipulate the mass set in motion, we apply a wrist weight. We use a TurnTuri sports wrist weight of 1kg.

#### Experiment protocol

The experiment was coded in Python using functions from PsychoPy. The experiment was controlled via a Brainvision Button Box (Brain Products GmbH, Munich, Germany), which was also streaming its output to the data collection PC unit.

#### Video and kinematics

The participants are recorded via a videocamera (Logitech StreamCam), sampling at 60 frames per second. We used Mediapipe^9^, to track the skeleton and facial movements, which is implemented in Masked-piper which we also use for masking the videos^10^.

#### Surface ElectroMyoGraphy (EMG)

We measured sEMG using a wired BrainAmp ExG system (Brain Products GmbH, Munich, Germany). Disposable surface electrodes (Kendall 24mm Arbo H124SG) were used, and for each of the four muscle targets we had 3 (positive, negative, ground) electrodes (12 electrodes total). The sEMG system sampled at 2500 Hz (for post-processing filters see below). We prepare the skin surface for EMG application with a scrub gel (NuPrep) followed by a cotton ball swipe with alcohol (Podior 70%). Positive and negative electrodes were attached with a 15mm distance center to center. We applied electrodes for focal muscles which directly participate in the internal (pectoralis major) and external rotation (infraspinatus) of the humerus. We attached the electrodes for focal muscles ipsilaterally (relative to the dominant hand). We attached electrodes to the muscle belly of the clavicular head of the pectoralis major, with a ground electrode on the clavicle on the opposite side. We also applied electrodes for postural muscles which will likely anticipate and react to postural perturbations due to upper limb movements. Since these muscles should act in the opposite direction of the postural perturbation of the dominant hand, we attach electrodes contralaterally to the moving dominant hand. We attach electrodes to the rectus abdominus and to erector spinae muscle group (specifically, the iliocostalis romborum), with ground electrodes on the iliac crest on the opposite side.

#### Ground-reaction forces

We used an inhouse-built 1m2 balance board with vertical pressure sensors. The sensors were derived and remodified from the Wii-Balance board sensors. The sampling rate was 400Hz. The system was time-locked within millisecond accuracy and has a spatial accuracy of several sub-millimeters. A national instruments card, USB-62221 performed the A/D conversion and was connected via USB to the PC.

#### Acoustics

To ensure proper acoustic intensity measurements we used a headset microphone; MicroMic C520 (AKG, Inc.) headset condenser cardioid microphone sampling at 16Khz. The gain levels of the condenser power source were set by the hardware (and could not be changed).

#### Recording setup and synchronization

We use LabStreamLayer (see here) which provides a uniform interface for streaming different signals along a network, where a common time stamp for each signal ensures sub-millisecond synchronization. We used a Linux system to record and stream the microphone recordings. Additionally, a second PC collected video and streamed ground reaction forces, and EMG. A data collection PC collected the audio, ground reaction force, and EMG streams and stored the output in XDF format for efficient storing of multiple time-varying signals.

### Experimental procedure

Participants were admitted to the study based on exclusion criteria and signed informed consent. Participants took off their shoes and we proceeded with the body measurements while instructing the participant about the nature of the study. After body measurements, we applied the surface EMG.

Upon start of the experiment, participants take a standing position on the force platform. The experiment commences with calibration and practice trials. Then, for the practice trials, each movement was practiced with expiring and vocalization while performing the movement conditions, and the participant is introduced to wearing the wrist weight of 1kg. After practice trials, participants performed 80 blocked trials. For each (practice) trial participants were closely guided by the information on the monitor. Firstly, participants are shown the movement to be performed for the trial and have to prompt the experimenter that they are ready to continue. Participants were instructed to adopt the start position of the movement, which is a 90-degree elbow flexion, with either an externally rotated humerus (start position for internal rotation), or a non-rotated humerus with the wrist in front of the body (rest position for the other movement conditions).

For the no movement condition participants were asked to rest their arms alongside their bodies. Upon trial start, participants inhaled deeply with a timer counting down from 4 seconds. Subsequently, participants were asked to continuously vocalize with a schwa sound, or expire, with a screen appearing after 3 seconds to perform the movement with visual guidance to where the movements end position is so that participants are reminded of the movement. After an additional 4 seconds, the trial ends, which allowed for more than enough time to perform the movement and stabilize vocalization after the perturbation. Participants were explicitly instructed to keep their phonation as stable as possible during the different movement conditions.

### Preprocessing of the data streams

To reduce heart rate artefacts artifacts we apply a common method^5^ of high-pass filtering the signal at 30Hz using a zero-phase 4th order Butterworth filter. We then full-wave rectified the EMG signal and applied a zero-phase low-pass 4th order Butterworth filter at 20Hz (after padding). We normalized the EMG signals within participants before submitting to analyses.

We upsampled the balance board from 400Hz to 2500 Hertz. We then applied a zero-phase low-pass 20Hz 2nd order Butterworth filter to the padded signals. As a key measure for postural perturbation, we computed the change in 2D magnitude (L2 norm of the center of pressure x and y) in the center of pressure (hereinafter COPc). For acoustics, we extract the smoothed amplitude envelope (hereinafter envelope). For the envelope we apply a Hilbert transform to the waveform signal, then take the complex modulus to create a 1D timeseries, which is then resampled at 2500Hz, and smoothed with a 12Hz Hanning window. All signals were sampled at or upsampled to, 2500 Hz (using linear interpolation).

### Overview data analyses

to be related to arm movement related muscle activity and postural activity. We determined amplitude deviation by first detrending the amplitude envelope timeseries, to express positive or negative peaks relative to this trend line (amplitude generally decreases when the lungs deflate).

We will measure the global maximum happening within the analyses window for the envelope, muscle activity, and the changes in the center of pressure. We will only analyze vocalization trials (and ignore expiration trials for this report). An example time series is shown for a flexion movement condition (Figure 2). At time = 0, the prompt is given to the participant to vocalize. We determine a detrending line using linear regression for the 1 to 5 seconds after the vocalization prompt. Note, At 3 seconds (3000ms) there is a movement prompt. However, we determine our window where we assess peaks in signals at 500 milliseconds before and after the movement onset/offset (using the peak finding function on the 2D speed time series of the wrist). In these trials, the analyses window is given in grey dashed bars, which is 500 milliseconds after and before movement onset/offset.

**Figure 2:**
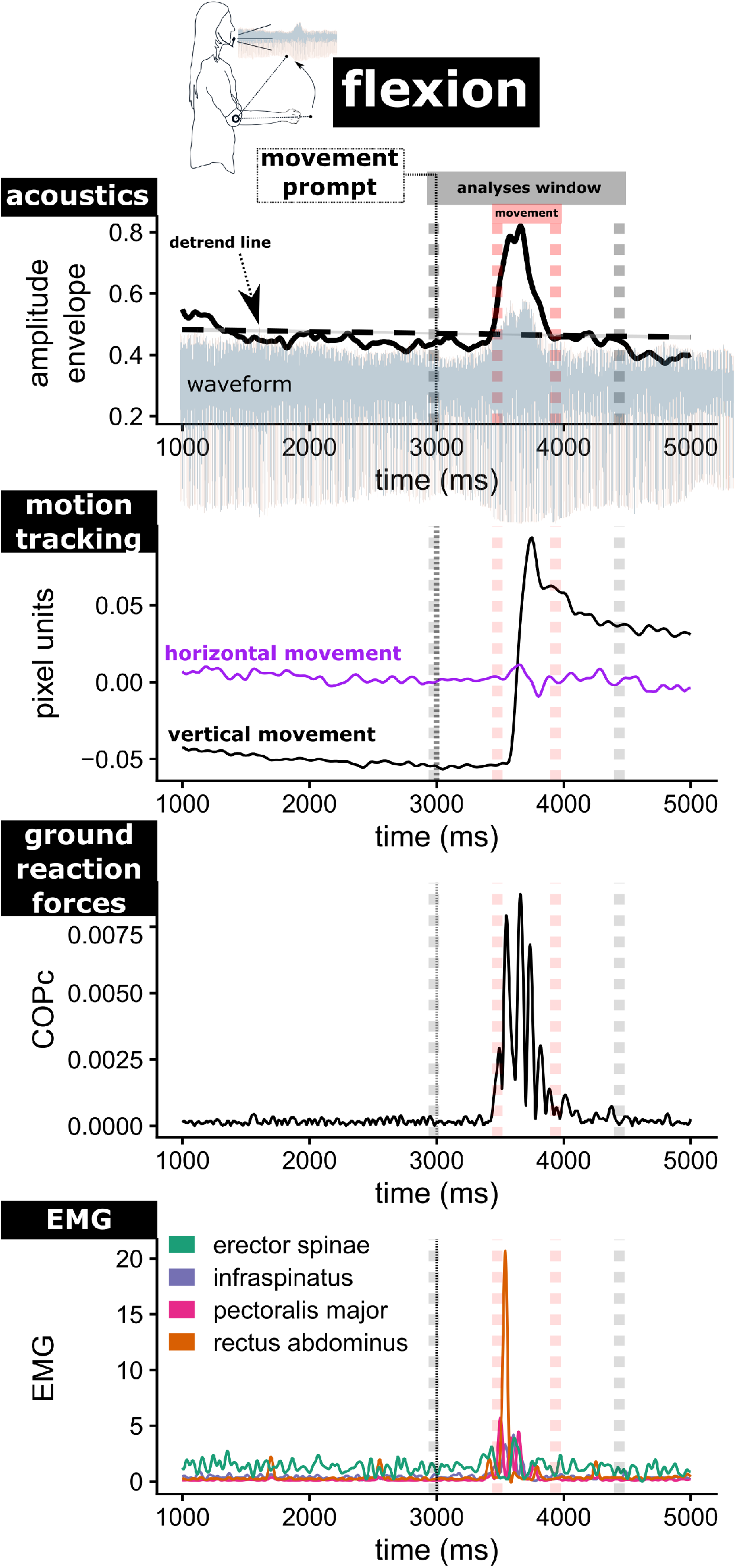
Time series example for a flexion movement + vocalization trial. *Note*. An example trial and their associated signals are shown (a flexion movement condition without wrist-weight). At time = 0 the prompt is given to the participant to vocalize. We determine a detrending line using linear regression for the 1 to 5 seconds after the vocalization prompt. Note, At 3 seconds (3000ms) there is a movement prompt. However, we determine our window where we assess peaks in signals at 500 milliseconds before and after the movement onset/offset (using peakfinding function on the 2D speed time series of the wrist). In these trials, the analyses window is given in grey dashed bars, which is 500 millseconds after and before movement onset/offset.

## RESULTS

Figure 3 provides an overview of the muscle activity patterns for each movement condition, split over weight conditions. The descriptive results suggest indeed that the different movement conditions solicit different kind of muscular activity patterns.

**Figure 3:**
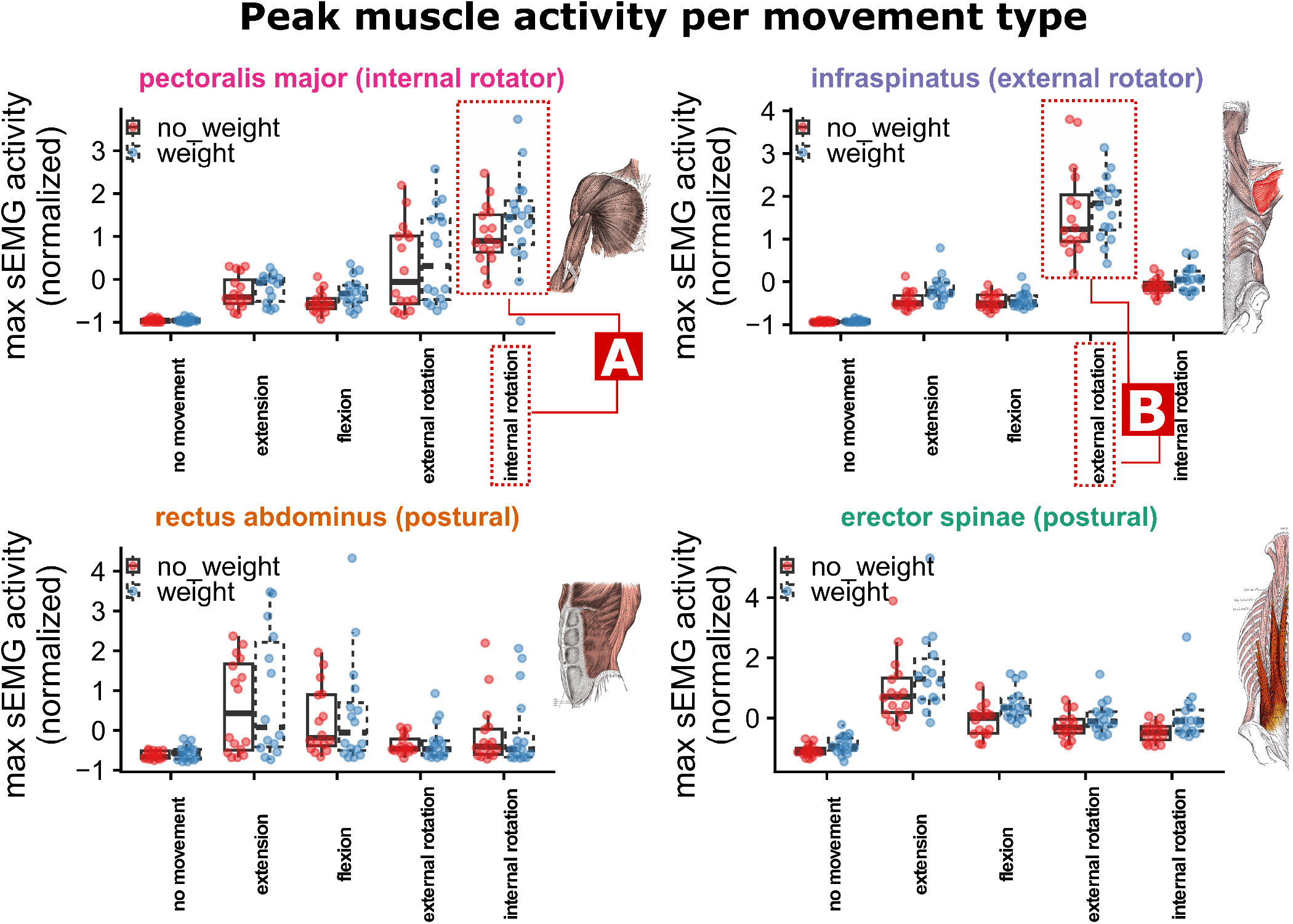
Time series example for a flexion movement + vocalization trial. *Note*. The peak muscle activity normalized for each muscle, is shown per movement and weight condition. A) highlights the fact that the pectoralis major, is - as to be expected - most active for the internal rotation movement condition. B) Also, to be expected, the infraspinatus is most active during the external rotation. Note further, that both postural muscles seem more active during extension (and secondarily flexion).

Figure 3 serving as a manipulation check, we can hereby ask three questions: 1) Do different upper limb movements lead to positively peaked deviations of the amplitude envelope of ongoing voicing? 2) Does peak muscle activity predict positively peaked deviations of the amplitude envelope of ongoing voicing, and 3) what muscle activity is related to posture-related dynamics?

### Effects of different movements on positive peaks in vocalization

We first modeled with a mixed linear regression the variation in positive peaks in the amplitude envelope (using R-package lme4), with participant as random intercept (for more complex random slope models did not converge). A model with weight and movement conditions explained more variance than a base model predicting the overall mean, Change in 2 (5) = 53.68, *p* < .001). The model coefficients are given in Table 1. It can be seen that there is a positive but not statistically reliable effect of wrist weight in this sample. Further, all movements (extension, flexion, internal rotation, external rotation) lead to statistically reliable increases in positive peaks in the amplitude envelope relative to the no movement condition (with flexion and external rotation leading to more extreme effects).

**Table 1:** Effects of weight and movement condition on positive peaks in the amplitude envelope.

| Predictor |  |  |  |  |
| --- | --- | --- | --- | --- |
|  | Estimate | SE | t-value | p-value |
| Intercept (no weight, no movement) | 0.164 | 0.101 | 1.63 | .181 |
| vs. weight | 0.062 | 0.063 | 0.98 | .328 |
| vs. extension | 0.436 | 0.100 | 4.35 | <.001 |
| vs. flexion | 0.618 | 0.100 | 6.17 | <.001 |
| vs. internal r. | 0.568 | 0.100 | 5.67 | <.001 |
| vs. external r. | 0.794 | 0.100 | 7.92 | <.001 |

### Effects of muscle activity on positive peaks in vocalization

We also directly relate muscle activity peaks with the positive peaks in the amplitude envelope. We first assessed the VIFs between the muscle activity peaks, which yielded a maximum VIF value of 2.91, and therefore we can combine the different muscle activity measurements in one model to predict amplitude envelope peaks. In a model with the participant as random intercept (a more complex random slope model did not converge) the different peak muscle activities explained more variance than a base model predicting the overall mean, Change in 2 (4) = 57.72, *p* < .001). The model coefficients are given in Table 2 and show that peak EMG activity in all the muscles (but especially the rectus abdominus, a well-known expiratory muscle) leads to statistically reliable increases in positive peaks in the amplitude envelope.

**Table 2:**
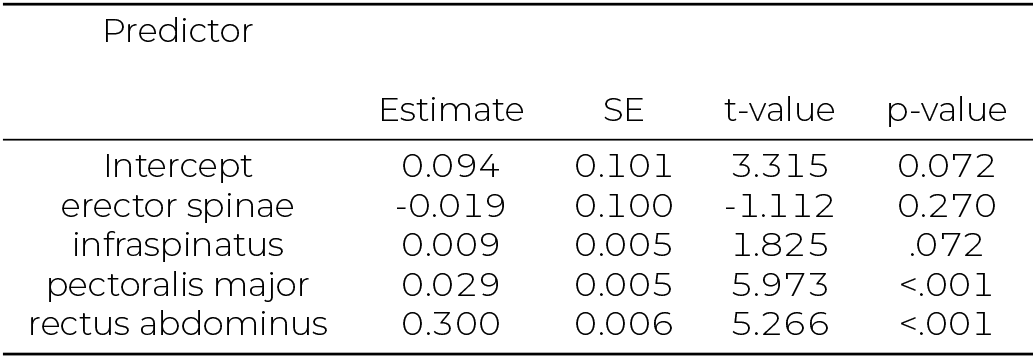
Effects of weight and movement condition on positive peaks in the amplitude envelope.

### Associations of muscle activity with changes in the center of pressure (COPc)

We similarly performed a linear mixed regression (with the participant as random intercept) with a model containing peak EMG activity for each muscle which was regressed on the peak in change in the center of pressure (COPc). We obtained that a base model predicting the overall mean of COPc was outperformed relative to said model, Change in 2 (4.00) = 64.22, *p* < .001). Table 3 provides the coefficient information. We find that only the postural muscles (rectus abdominus, erector spinae) indeed reliably predict the magnitude of changes in the center of pressure, while the pectoralis major and the infraspinatus do not reliably relate to the changes in ground reaction forces.

**Table 3:** Effects of muscle activity on change in center of pressure.

| Predictor | Estimate | SE | t-value | p-value |
| --- | --- | --- | --- | --- |
| Intercept | -0.886 | 0.290 | -3.051 | 0.146 |
| erector spinae | 0.112 | 0.035 | 3.166 | 0.002 |
| infraspinatus | 0.003 | 0.011 | 0.260 | 0.796 |
| pectoralis major | 0.016 | 0.010 | 1.613 | 0.111 |
| rectus abdominus | 0.074 | 0.012 | 6.219 | <.001 |

**Figure 4:**
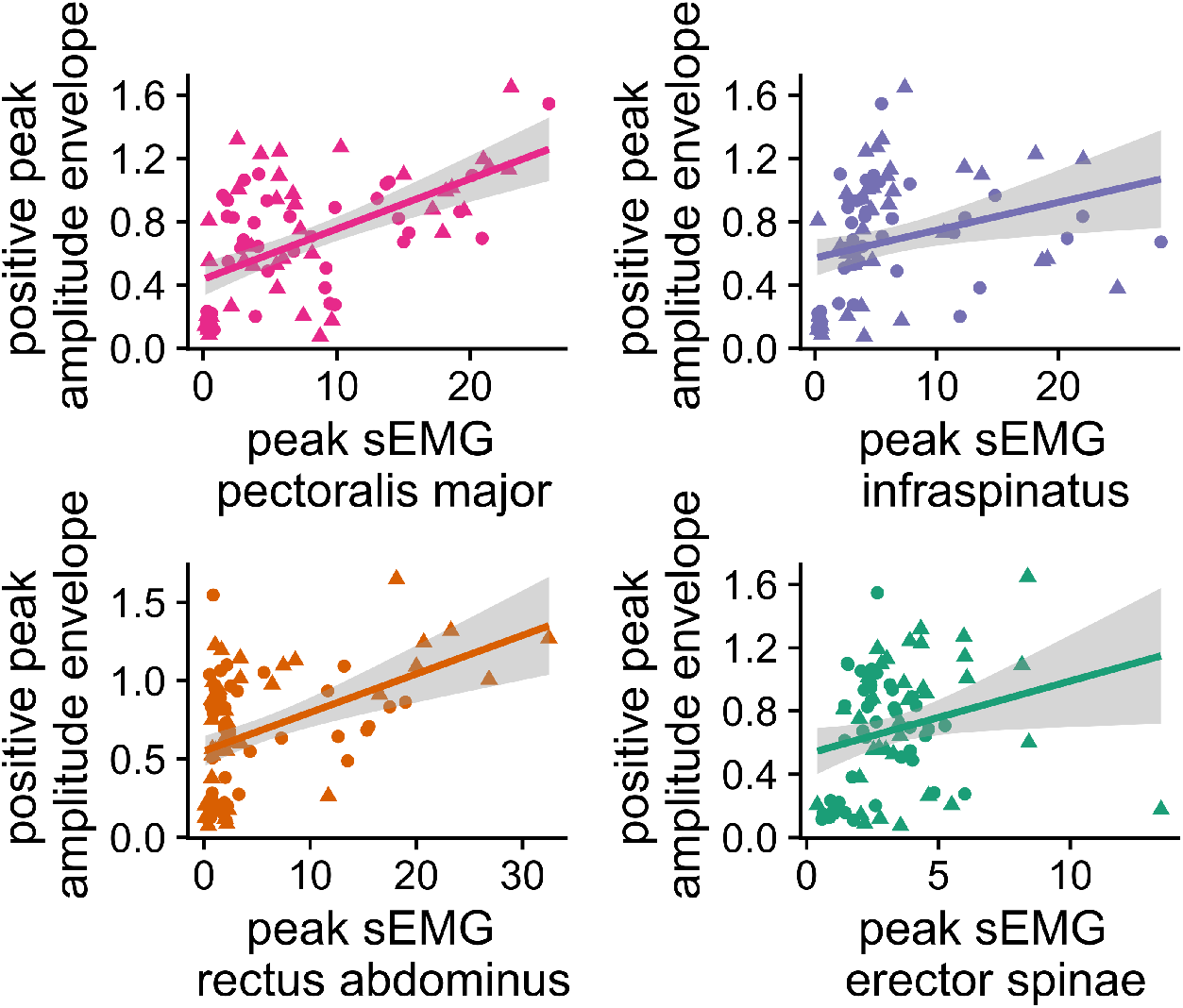
Relations between peak muscle activity and positive peaks in the amplitude. *Note*. Triangles indicate movements with wrist-weight.

## DISCUSSION

In summary: Movement versus non-movement of the upper limbs yields unintentional positive peaks on the amplitude envelope of vocalization. This can be labeled as ‘unintentional’ as the task is to produce a stable vocalization output. Further, we show that particular muscle activity is reliably related to positive peaks in the voice amplitude, especially the rectus abdominus, an expiratory-associated muscle. This muscle and the erector spinae are further also found to be related to postural stability, as the change in the center of pressure reliably positively related to the these postural muscle activations (in contrast to the other focal muscles). We observe small but statistically unreliable effects of a 1kg wrist weight has an effect onto the amplitude envelope. With a confirmatory study we will be able to confirm this effect of wrist weight, which would confirm a role of force-transfers as the mass of the moving segment is increased, therefore yielding more force per unit acceleration ^14^. From a more detailed study of the data than reported here, we will be able to ascertain whether certain muscles can also affect the voice by decreasing subglottal pressure (thereby decreasing amplitude). Once this research has been performed in a more detailed way, we will be able to make clear predictions about what type upper limb movements have a certain effect on the voice, which may then help understand how gestures may be functionally integrated with the prosodic targets dictated by the particular prosodic regime of a spoken language ^12^.

## CONCLUSION

These results show that the voice is affected by posture-affecting muscle activity induced by upper limb movement. We thereby show that voice production is a dynamically open system, that will be affected by other communicative actions such as hand gesture. This has deep implications for why gesture and speech are often synchronized on the prosodic level [5]. This study thereby confirms that we should not forget that the acoustic signal is produced by a complex set of interactive processes that relate to respiratory control. Understanding such processes will undoubtedly better tune speech-recognition and synthesis technology.

**Figure 5:**
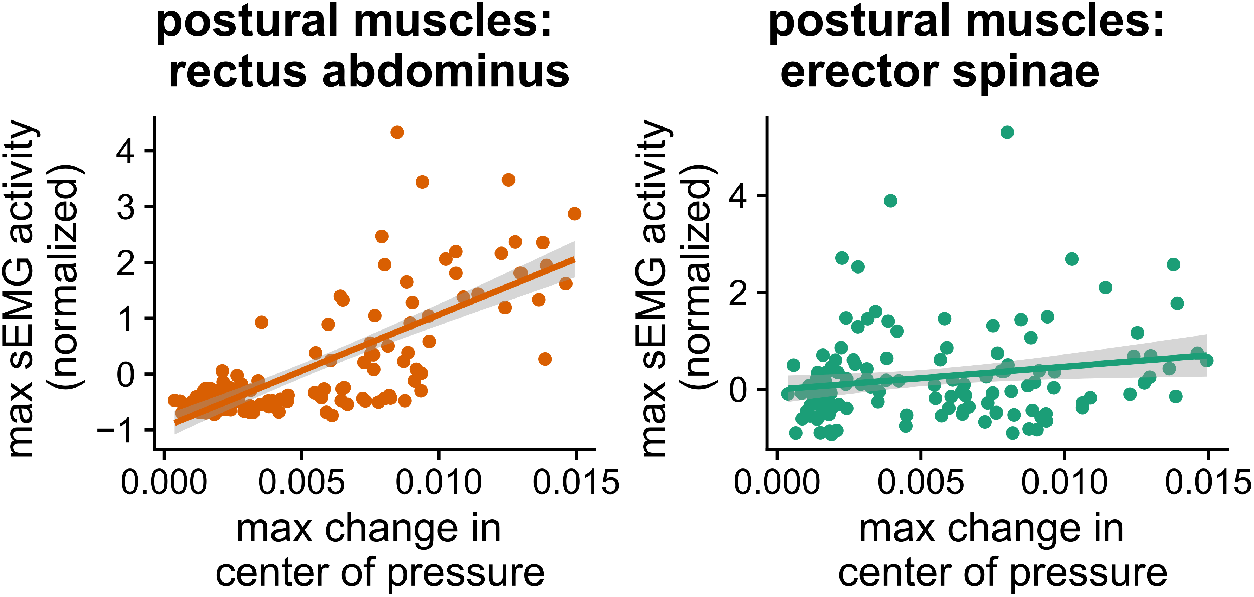
Relations between peak muscle activity and magnitude change in the center of pressure.

## ACKNOWLEDGMENTS

This research has been funded by a VENI grant (VI.Veni 0.201G.047: PI Wim Pouw) awarded by the Dutch Research Council (NWO). We would like to thank Pascal de Water from the Donders Institute for his technical support for creating the LabStreamingLayer capabilities that are used in this research, and for his general guidance that has tremendously improved this research.

## DATA AVAILABILITY

The data supporting this preprint will be opened alongside the publication of a more detailed pre-registration.

